# Education in the Time of COVID-19: The Improvised Experiment of Virtual Assessments

**DOI:** 10.1101/2022.11.01.514696

**Authors:** Esteban Guevara Hidalgo

## Abstract

One of the aspects in which the COVID-19 pandemic impacted the most was education. Teachers and students had to face a new reality for which they were not prepared adapting in an improvised way new methods and strategies to teach and to learn. Within virtual education, exams reduced in some cases to multiple choice tests while others tried to mimic traditional (pen and paper) exams. In this paper, these two kind of evaluations are compared. Although the results appear to be similar, a deeper look shows that their structure is completely different and some groups of students are unfairly harmed or benefited depending on the assessment applied. Beyond analyzing the reasons of this discrepancy, it is determined that for some type of evaluation, at least 21.1% of students maybe passing a course irregularly meanwhile, at least 5.5% could be failing a course despite their actual capabilities.

## 1. Introduction

Academic integrity [1–8] is the cornerstone of education. Educational institutions have embedded this principle in their constitutions as codes of honor [5,7,9], establishing, in some cases, strong punishments to the offenders, as without it, the educational process lose all its sense. Cheating has always been a concern [1,10–12], and different control mechanisms have been perfected during the time in order to reduce the occurrence of such events [13], specially during exams season. These mechanisms vary depending on the institution and the region, and, in occasions, they can be very strict, regarding the stuff a student can bring to an exam, the dress code, the distance among students, the looks, etc.

However, academic dishonesty has accentuated worryingly during the COVID-19 pandemic due to virtual education [14–23]. Many efforts have been made attempting to translate the “non-virtual” control mechanisms to a virtual environment [19,17,24–27] using platforms and technologies addressed in this direction [19]. However, because of their cost, the large majority used simpler alternatives, where results more difficult to (successfully) avoid cheating. On the other hand, evaluations have reduced in most cases, only to multiple choice questions (MCQs) tests where the answers can be easily shared in one of the several social networks or instant messaging apps favoring the cheating [28–30]. In order to avoid this issue, others have tried to mimic the traditional (pen and paper) exams. These assessments were based on the development of an unpublished problem, and not only in its answer, putting aside from any platform, and specially from MCQ tests. However, despite these efforts to difficult dishonest behaviors, cheating has been still detected.

In this paper, we present the results obtained in a experiment over a group of students which were examined with these two kind of assessments. Although the average outcome was similar independently of the type of evaluation, the distribution of grades among students changed, i.e., some students performed better than others depending on the type of exam. Moreover, the students who obtained the lowest grades in one kind of test improved (irregularly) in the other. Meanwhile, the students who obtained the higher grades diminished their performance. Apart from analyzing the causes of these discrepancies, three scenarios are presented and it is shown for which the best results are obtained, and which group of students is unfairly benefited.

### 1.1. Education in Pandemic Time

The core problems in education are still discussed as they evolve. The discussion goes from the method used to teach to the proper way of evaluating the knowledge of a student. Although these concerns have been answered in several occasions through the years, there is not an universal answer. However, even before the pandemic, methods which have been proved to be ineffective and obsolete [31], continued being used protected by the so called “academic freedom”. The “new education” identified the flaws of these methods and moved in the direction of the new technologies [32]. This inclusion was a necessity to face new challenges like lack of interest, short attention spam, and a society fully absorbed in that same technology. So as the use of tablets, computers, smartphones, messaging apps, social networks, and YouTube have become a daily basis aspect of our lives, it is almost impossible to conceive education without them [33]. Nonetheless, adopting them correctly was a reality only for the few which went step by step in that direction. For the large majority, the transition to virtual education was forced and abrupt due to COVID-19.

#### 1.1.1. Teaching in Pandemic Time

With some exceptions, in most of the cases, virtual education was reduced to the presentation (of a large amount) of PowerPoint slides [31,34]. In some areas like social sciences, this always has been the case, but during the pandemic, many important activities in engineering and science like labs and exercises were also boarded in this way. If we add to this the sometimes completely lack of interaction between the students and the teacher, the class became in nothing but a monologue of the teacher reading and passing slides. Some efforts were made trying to incorporate (in an improvised way) new strategies which have appeared in recent years in a virtual and non-virtual education context. For example, the flipped classroom [35–37] and gamification [38–42] introducing within the lectures: videos, presentations, crosswords, quizzes, quests, short tests, among others. However their objective has been distorted resulting in nicer slides, poor quality lecture videos but specially, student work overload.

Within the flipped classroom, either the teacher has recorded himself giving a lecture so they can discuss it and work in examples and projects during the synchronous course or he has assigned the students videos already available with the same purpose. However, although flipped classroom has been implemented in non-virtual environments with successful results [36,37], one of its major advantages has been wasted, which is having more available time to interact with the student. In practice, if this method is implemented by most of the teachers, apart from homework, projects and other additional activities, the students have to expend an extra time to watch the lecture videos resulting in a work overload [35].

In some other cases, the lectures adopted a precarious MOOC (Massive Open Online Course) structure [43–45]. The material and the course activities were loaded to a platform where the students can access them whenever they want. The material generally consists in the slides of the course, sometimes videos, and some additional material like extended readings. The course was generally graded via projects, crosswords, games, puzzles, and MCQ tests. All this, with almost no contact with the instructor or any control mechanism.

The possibility of reviewing a topic as many times as needed and at ones own pace is an advantage which, in general, was not present before the pandemic. However, due to these videos and material are available after the lectures, attending them has lost their purpose, specially when the content of the material is exactly the same as the one “explained” by the teacher during the synchronous lectures [34]. The challenge, of course, is big as entire courses available in YouTube and other platforms are in fact better than the actual lectures [46]. For this reason, the teacher should use all these new methods, resources and tools to attract and keep the attention and interest of the student either to complement or to introduce him to a topic giving the student a reason to attend lectures beyond a test.

Far off the pros and cons of what was presented above and others not mentioned here, they all share the same problems which have been left unattended: academic dishonesty, student work overload, and lack of interaction (teacher-student or student-student).

#### 1.1.2. Evaluations in Pandemic Time

The course score has been distributed among several activities like short tests, projects, homework, videos, posters, and other shorter ones like games, quizzes, crosswords, quests, etc. However, when this is repeated in most or every course, the result is an unmotivated, frustrated and stressed student overloaded with daily tests and pendent activities. On the other hand, exams have been reduced to MCQ tests (**v-Tests**). During the evaluation, a number of problems are loaded “randomly” from a question bank. The student either has to choice between several options, or to enter the answer into a box. The procedure is not evaluated, only the answer. It is not know how that answer was obtained and there is no difference between not knowing something at all and making small arithmetic mistakes. Alternatively, some teachers had mimic the traditional (pen and paper) exams into a virtual environment. These evaluations are based in the development of a problem (e.g., an exercise), where not only the answer is graded but also the process (**p-Tests**). Some control mechanisms can be implemented during these virtual tests to guaranty a honest behavior. For example, the use of the camera/s or the sharing of the screen. However, in the case of v-tests they are highly limited for the easiness of the sharing of an answer, specially when the camera only focus on the face of the student and it is not possible to see his desktop (and what is on it) and the tabs open in the computer. Apart from this concern, there are some issues that arise in the design of evaluations related to the size of the question bank or if the questions are modified versions of already available problems or they are unpublished.

## 2. Methodology

The study was performed on a group of 109 students from the engineering first year taking **Course X** in the fall semester 2021 during the COVID-19 pandemic. The lectures were carried out “virtually” using an electronic pen on a note-taking program. The group was created gathering students from three classrooms from 6 very different engineering careers: class A: 41 students, class B: 32 students, and class C: 36 students that were assigned to a same teacher at the beginning of the semester. The first part of the semester (*B*_1_) the students were evaluated using two **p-tests** and the second part (*B*_2_) with two **v-tests**. Additionally, there was a control (in the form of a p-test) during *B*_2_ in between the two v-tests. The study did not require a review and approbation of the institution ethics committee nor the participant consent as it consists in a direct analysis of the data of what actually occurred during the fall semester 2021.

### 2.1. The Tests Types and the Settings

#### 2.1.1. Written Procedure Virtual Tests: p-tests

Each p-test had a duration of 1 hour and consisted in four unpublished problems. The students were instructed to the access the meeting 15 minutes before the exam, to avoid problems like software updates, computer reboot, etc. Later, the teacher proceeded to check the student location which was indicated previously (both during the lectures and also by email). The camera should focus not only their faces but also their screens, hands and desks. This revision took an average of 30 minutes after which, the exam was projected/shared on the student’s screens. Once the exam started the students were not allowed to used the keyboard, mouse or smartphone. They had to solve the four problems “by hand” on a sheet of paper, and one hour later, scan it with some phone app and send it to the teacher’s email and also to load it to the platform. This process took an average of 5 minutes but the students were given 10 minutes to submit the exam’s pdf. The tests were graded using an electronic pen considering the procedure and not only the final answer, so small arithmetic mistakes did not affected greatly the final grade. The p-tests settings were designed entirely by the teacher.

#### 2.1.2. Multiple Choice Virtual Tests: v-tests

Each v-test had a duration of 50 minutes and consisted in five multiple choice problems. For each student, an exam was generated randomly by the platform from a database of around **20-30 exercises**, each one with five possible answers. The database was created with contributions of the 7 course teachers. The problems not necessarily had to be unpublished but modifications of exercises from the homework sheets. The configuration of the exam in the platform was so the difficulty be the same for every student. The v-tests were applied to 795 students, (the total number of students taking Course X the fall semester 2021) including the 109 students that were part of this study and that were evaluated using p-tests during *B*_1_. To avoid network and platform issues, the students were separated in two groups, the first half took the exam at a determined time and one hour later the second group, both from the exact same database. Although the students were told (by email before the exam) to locate as for the p-tests, due to the v-tests settings, a location control was not possible before but during the exams. However, this was not a rule, but an arbitrary decision of each teacher. Meaning that many of the 795 students took these v-tests with a minimum control (with the camera focusing only their faces). The students were not allowed to use the keyboard, mouse, smartphone or notes, however this was not necessarily possible to verify. The exam was automatically graded by the platform considering only the final answer and not the process or how that answer was obtained. Small arithmetic mistakes could affect drastically the final outcome. The teacher had not control over the v-tests settings which were designed by the course chair.

#### 2.1.3. p-Grades and v-Grades

The p-grade is the average of the two p-tests grades taken during *B*_1_ and the control taken during *B*_2_. The v-grade is the average of the two v-tests grades (during *B*_2_). Both the p- and v-grade over 10 points. The final grade is the sum of the p-grade and the v-grade. If the final grade is less than 5*/*20 points the student (directly) fails the course, if it is greater o equal than 10*/*20 points, passes it, and if it is in between, it is necessary to take an additional exam. Lab and class activities are also considered in the student score (in fact, they represent the final’s grade 40%). However, it is important to remark, the analysis presented in this paper compares only the tests performance. Thus, the grades and their scale for passing or failing are different from the actual ones.

## 3. Results

### 3.1. Students Redistribution during the v-Tests

The p-grades were considered as reference values for the following analysis. The results ranges from 0 to 9 points while for the v-grades from 0 to 7.5, (both over 10). The fact that v-grades can only take specific values is evidenced as stairs in the data. The p- and v-grades sorted in ascending order are shown in Fig. 1. Depending on the p-grades performance, the students were classified in three groups: *G*_1_ corresponds to the 1/3 of students with the lowest performance (with grades from 0 to 2.22). *G*_2_, students with grades from 2.22 to 5.475 and, *G*_3_ is the top 1/3 (Fig. 1(a)). Although the average performance is similar independently of the type of evaluation 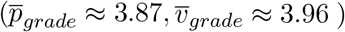, there is a change in the performance of the students during the v-tests, and thus, in their distribution within the groups. This students redistribution is shown in Fig. 1(b) using the same colors as for the group classification made from the p-tests results.

**Figure 1.**
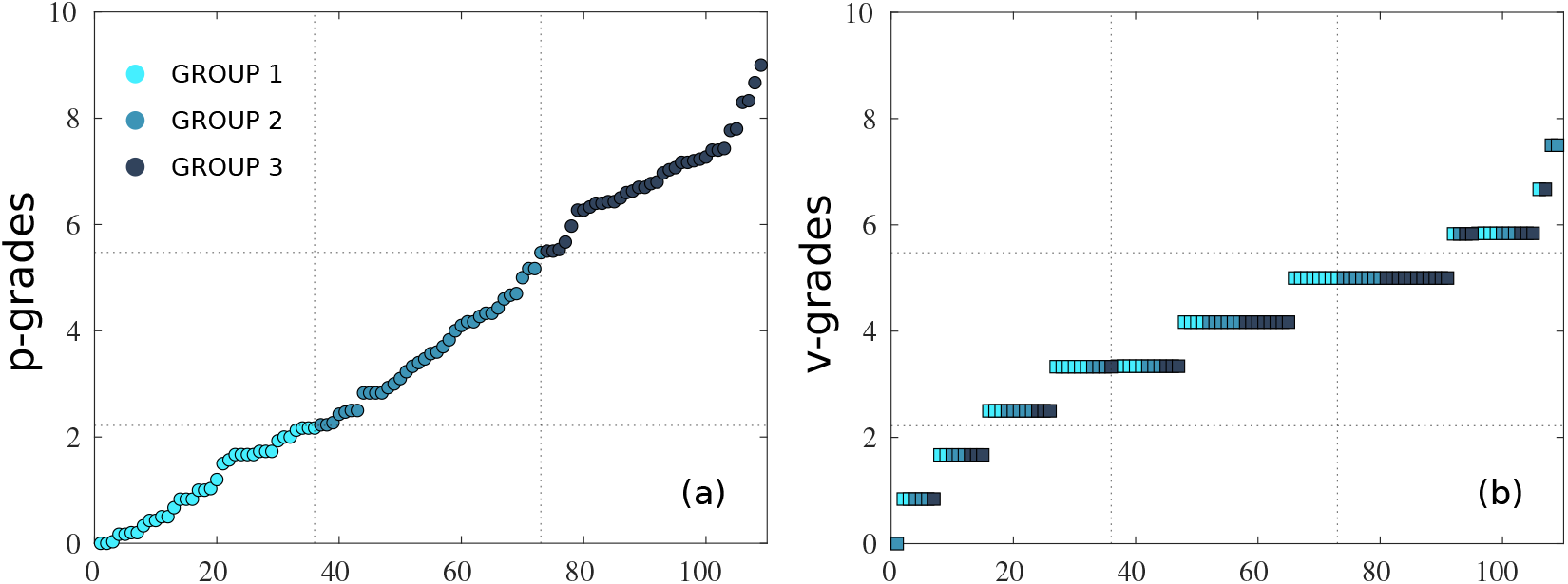
**(a)** Average of the three p-tests results, two tests were taken during the first part of the semester, and a control during the second part in between the two v-tests whose average is presented in **(b)**. The data is presented in ascending order for each of the 109 students. The students were classified in three groups (based on the p-grades results): *G*_1_, the bottom 1/3, *G*_2_, the middle group, and *G*_3_, the top 1/3. How the students performed and redistributed during the v-tests is shown in (b) using the same group classification as for (a).

In a closer look, students who obtained the lowest grades during the p-tests, improved considerably in the multiple choice tests. Meanwhile, the students who obtained the higher p-tests grades diminished their performance. Specifically, students from *G*_1_ improved an average of 2.83 points while students from *G*_3_ worsened and average of 2.71 points (Fig. 2). This last result is expected, as the procedure is not graded affecting directly the final outcome, however the first result is striking. Except from 2 (of 36) *G*_1_-students, the rest improved their performance between 0.34 and 6.24 points during the v-tests (Fig. 2 (a)). The *G*_2_-behavior is diverse, some students got better (17/37) and others got worse (20/37), but the average performance of the group remains approximately the same (Fig. 2 (b)). Most *G*_3_-students (excepting for 1 of 36), worsen their performance between 0.03 and 6.33 points during the v-tests (Fig. 2 (c)).

**Figure 2.**
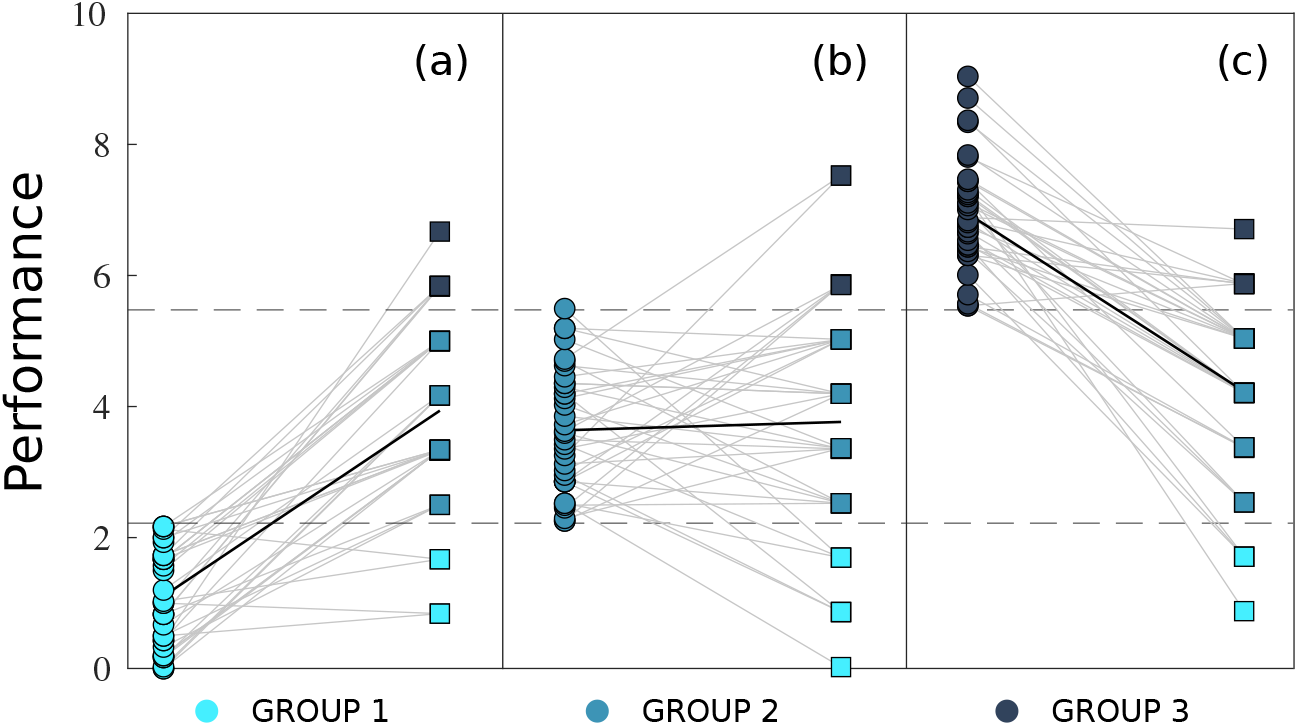
Groups performance during the v-tests: *G*_1_-students improved an average of 2.83 points, and up to 6.24 points during the multiple choice virtual tests (v-tests) applied during *B*_2_ (a), while *G*_3_-students diminish their performance an average of 2.71 points, and up to 6.33 points (c).

### 3.2. Passing Scenarios

Apart from the original **Scenario 1 (M)** described above (Fig. 3(a)), other two hypothetical scenarios are also considered assuming the students were evaluated only with p-tests: **Scenario 2 (P)** (Fig. 3(b)), and only with v-tests: **Scenario 3 (V)** (Fig. 3(c)). These last two scenarios were created from the actual v-and p-grades. Figure 3 presents the final grades *X* (over 20) in ascending order for every student in each scenario. The colors correspond to the original group classification as in Sec. 3.1. The vertical dash lines define three zones which determine if a student fails the course (*Z*_1_: *X* < 5), has to take an additional exam (*Z*_2_ : 5 ≤ *X* < 10) or passes the course (*Z*_3_ : *X* ≥ 10).

**Figure 3.**
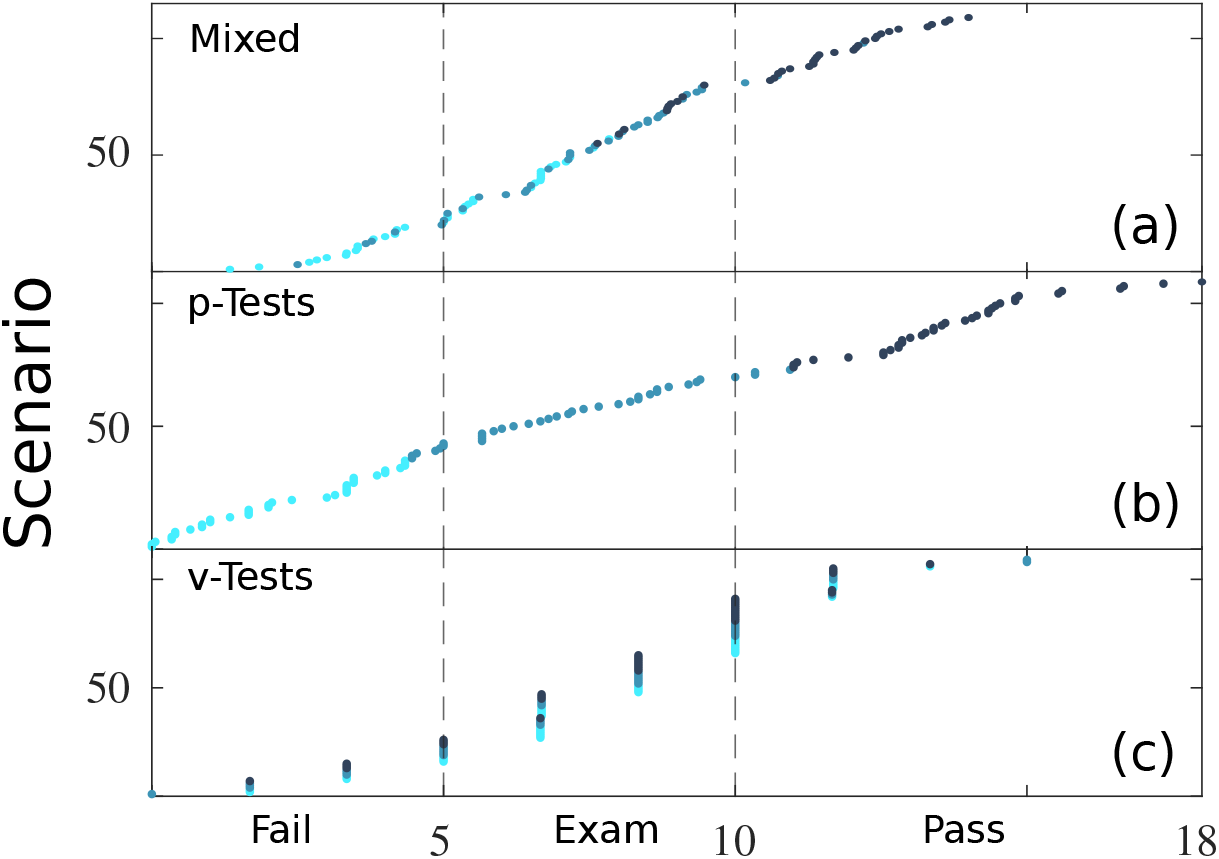
Final grades (over 20) in ascending order for every student in three scenarios. (a) Scenario 1 (M): the student was evaluated with p-tests and v-tests. (b) Scenario 2 (P): only with p-tests and (c) Scenario 3 (V): only with v-tests. The vertical dash lines define if a student fails the course (Zone 1), has to take an additional exam (Zone 2) or passes the course (Zone 3). The colors correspond to the original group classification given in Sec. 3.1. The group and zone composition, and the passing percentages are presented in Tables 1, 2, and 3.

Although the average final grade does not presents major differences 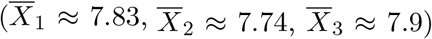 between scenarios, the zone composition does. While for Scenario 1 (M) and Scenario 2 (P), *Z*_1_ and *Z*_3_ are composed (as expected) mainly from students from *G*_1_ and *G*_3_ respectively (Table 1 and 2), for Scenario 3 (V) the performance structure is broken, students from every group are distributed across every zone apparently without any relation (Table 3).

**Table 1.**
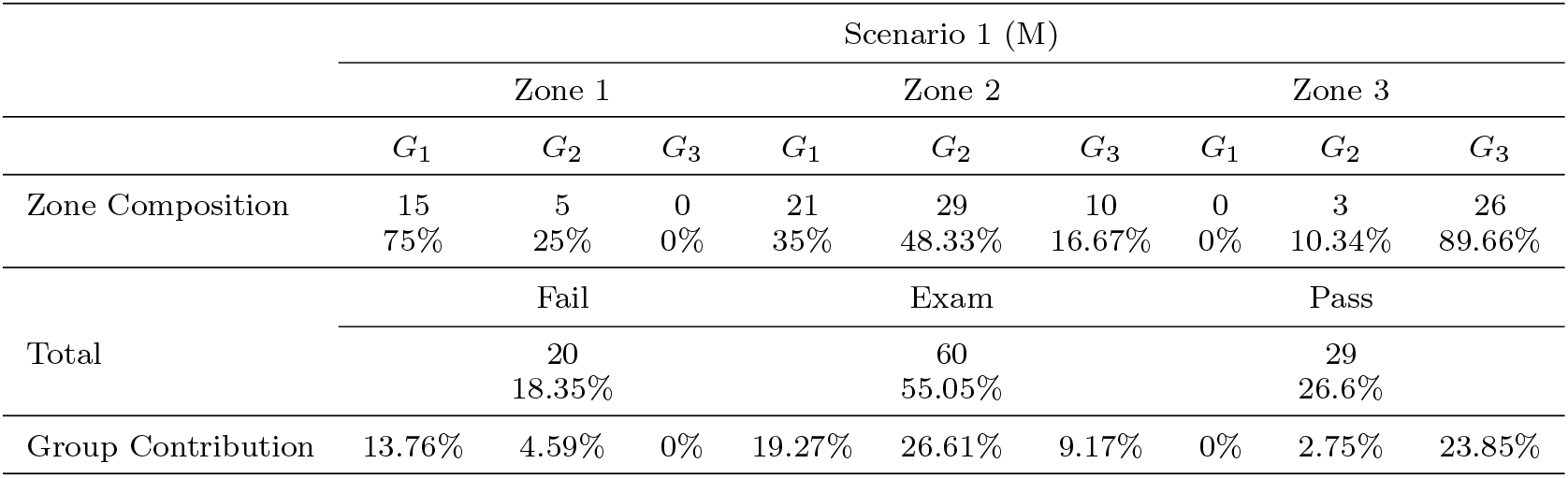
Scenario 1 (M): p-tests and v-tests (Figure 3(a)).

|  | Scenario 1 (M) |  |  |  |  |  |  |  |  |
| --- | --- | --- | --- | --- | --- | --- | --- | --- | --- |
|  | Zone 1 |  |  | Zone 2 |  |  | Zone 3 |  |  |
| | $G_1$ | $G_2$ | $G_3$ | $G_1$ | $G_2$ | $G_3$ | $G_1$ | $G_2$ | $G_3$ |
| Zone Composition | 15<br>75% | 5<br>25% | 0<br>0% | 21<br>35% | 29<br>48.33% | 10<br>16.67% | 0<br>0% | 3<br>10.34% | 26<br>89.66% |
| Total | Fail |  |  | Exam |  |  | Pass |  |  |
|  | 20<br>18.35% |  |  | 60<br>55.05% |  |  | 29<br>26.6% |  |  |
| Group Contribution | 13.76% | 4.59% | 0% | 19.27% | 26.61% | 9.17% | 0% | 2.75% | 23.85% |

**Table 2.** Scenario 2 (P): Only p-tests (Figure 3(b)).

|  | Scenario 2 (P) |  |  |  |  |  |  |  |  |
| --- | --- | --- | --- | --- | --- | --- | --- | --- | --- |
|  | Zone 1 |  |  | Zone 2 |  |  | Zone 3 |  |  |
| | $G_1$ | $G_2$ | $G_3$ | $G_1$ | $G_2$ | $G_3$ | $G_1$ | $G_2$ | $G_3$ |
| Zone Composition | 36<br>87.8% | 5<br>12.2% | 0<br>0% | 0<br>0% | 28<br>100% | 0<br>0% | 0<br>0% | 4<br>10% | 36<br>90% |
| Total | Fail |  |  | Exam |  |  | Pass |  |  |
|  | 41<br>37.61% |  |  | 28<br>25.69% |  |  | 40<br>36.7% |  |  |
| Group Contribution | 33.03% | 4.59% | 0% | 0% | 25.69% | 0% | 0% | 3.68% | 33.02% |

**Table 3.**
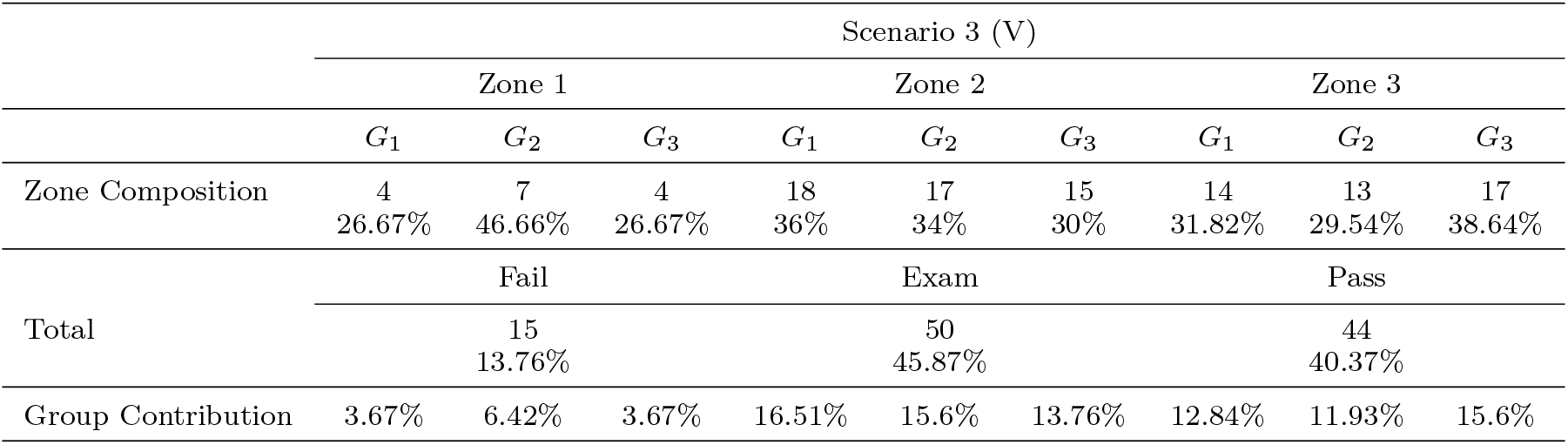
Scenario 3 (V): Only v-tests (Figure 3(c)).

Although the worsen of *G*_3_-students during v-tests could be related with small arithmetic mistakes affecting drastically their performance, it is important to consider the possibility that the *G*_1_-improvement could be related with the v-tests poor control mechanisms. Even worst, it is possible the v-tests results could be absolutely random and not related with the actual knowledge or performance of a student. Given that the best results (higher passing and lower losing percentage) are obtained in Scenario 3 (V) (40.37% direct passing and 13.76% direct failing)(Table 3), it is also possible some students under this Scenario could be failing or passing the course irregularly.

### 3.3. Beyond the Data

Despite previous works about proctoring in online exams [24,22], the group classification in Sec. 3.1, based on p-tests results, could be considered arbitrary. However, behind these results there is important hidden information. In every case, the p-tests results corresponded exactly with the student performance during the lectures. The best p-grades were obtained by students which attended the lectures, turned their cameras on, answered correctly questions, and showed interest. On the other hand, the worst grades were obtained by students whose participation was not only poor or null, but also by students which were sanctioned for cheating, by students which handed out an empty exam or did not attend the lectures at all. Thus, the *G*_1_-improvement becomes unlikely and suspicious, implying these multiple choice virtual tests (v-tests) could be favoring a wrong group of students and harming other.

Figure 4 compares the zone composition (from the data in Tables 2 and 3) of Scenario 3 (V) with respect to Scenario 2 (P). *n*_*P*_ is the number of students from a determined group in a determined zone from Scenario 2 (P), and similarly for *n*_*V*_. The variation Δ*n* = *n*_*V*_ − *n*_*P*_, measures the number of students who have moved from a zone *i* and group *j* in (P), 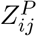, to another zone and group during the virtual scenario (V), 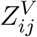. It is observed that 32 students from *G*_1_ in *Z*_1_ 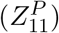 (the lowest p-tests performances) migrated to Zone 2 (18) and to Zone 3 (14), 11 students moved from 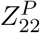 to Zone 1 (2) and to Zone 3 (9), and 19 students from 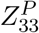 (the best p-tests results) migrated to Zone 1 (4) and to Zone 2 (15). Under Scenario 2 (V), before the additional exam, it is possible that at least a 21.1% (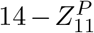 and 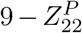) of the students maybe passing irregularly the course. This percentage increases to 25.47% after the additional exam. A 66.67% of students which maybe passing irregularly belongs to *G*_1_. On the other hand, at least 5.5% of students (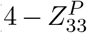 and 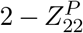) could be failing the course despite their actual capabilities. After the exam, this percentage increases to 11.92%, from which 84.62% are *G*_3_-students. Repeating a similar analysis for Scenario 1 (M), there is no migration to direct passing or failing, instead, 21 students from 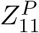 and 11 from Zone 3 have to take an additional exam 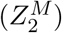. After the exam, only one 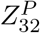-student could be failing the course unfairly and there is no apparent irregular passing.

**Figure 4.**
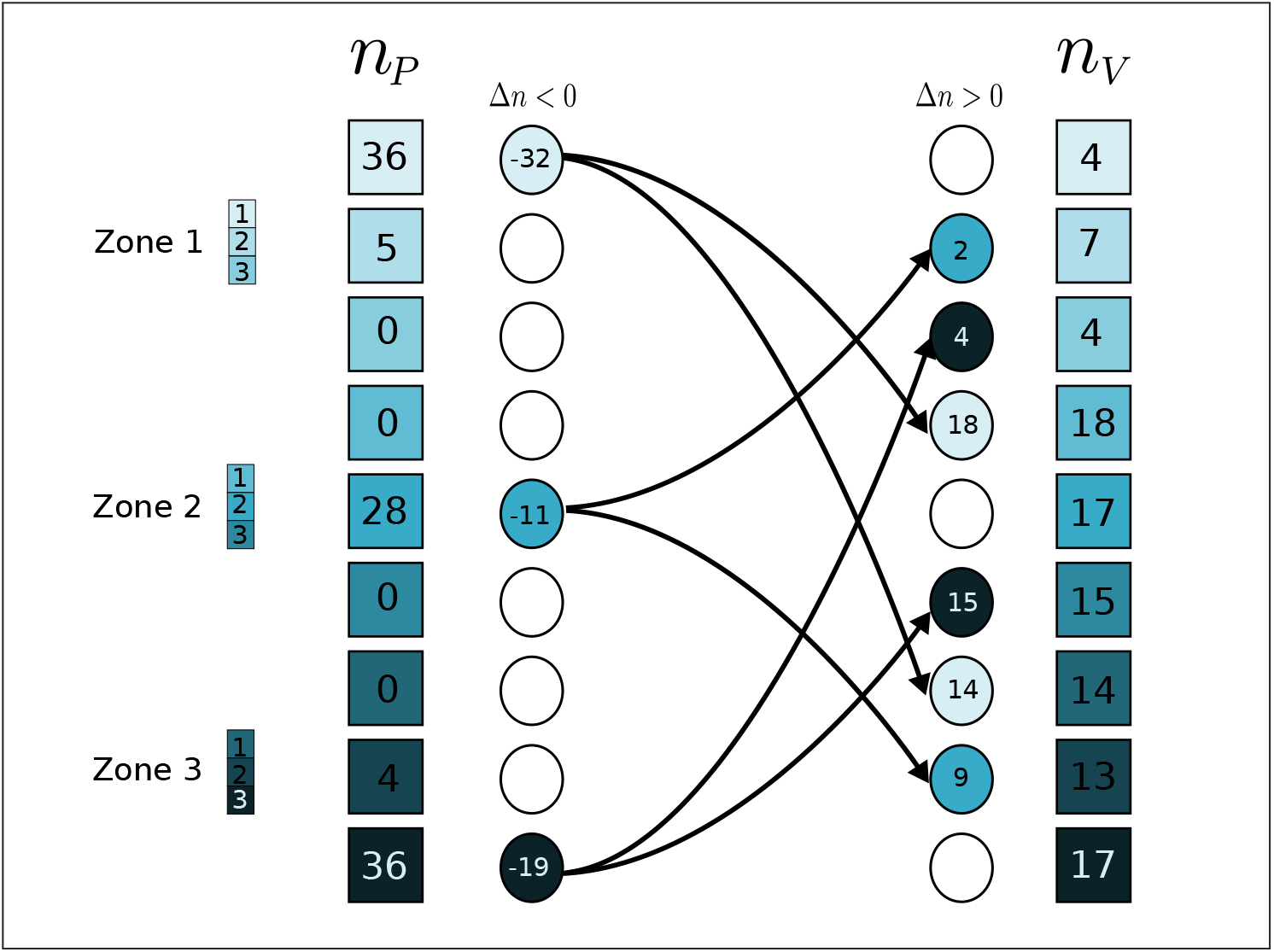
Comparison between the Zone Compositions of Scenario 2 (P) (left) and Scenario 3 (V) (right) (from the data in Tables 2 and 3). *n*_*P*_ is the number of students from a determined group in a determined zone from Scenario 2 (P), and similarly for *n*_*V*_. The variation Δ*n* = *n*_*V*_ − *n*_*P*_, measures the number of students who have moved from a zone *i* and group *j* in (P), 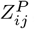, to another zone and group during the virtual scenario (V), 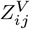. Before the additional exam, at least a 21.1% (14 students from 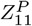 and 9 students from 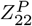) of the students maybe passing irregularly the course. On the other hand, at least 5.5% of students (4 from 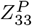 and 2 from 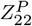) could be failing the course despite their actual capabilities. After the exam, these percentages increased to 25.47% (66.67% belongs to *G*_1_) and to 11.92%(84.62% are *G*_3_-students), respectively.

### 3.4. The Virtual Panacea

Before COVID-19, Course X evaluations were made via traditional exams. They were common to all first semester students and they were taken in auditoriums. During the pandemic, the course was evaluated entirely via v-tests (with the only exception of the students which participated in this study). An abrupt change in the passing percentage related to the transition to virtual education can be observed (Fig. 5). Although, a similar behavior is observed for all the first year courses, these differences in the passing percentage improvement is directly related with the teaching resources and with the way the course was evaluated. The higher improvement (35.87%) is observed for the course where the lectures were taught with PowerPoint slides and common v-tests and the lower improvement (6%) by a course managed with optical pencil and non-common p-tests. For the rest of courses, either each teacher decided the evaluation format, or they switched eventually to p-tests.

**Figure 5.**
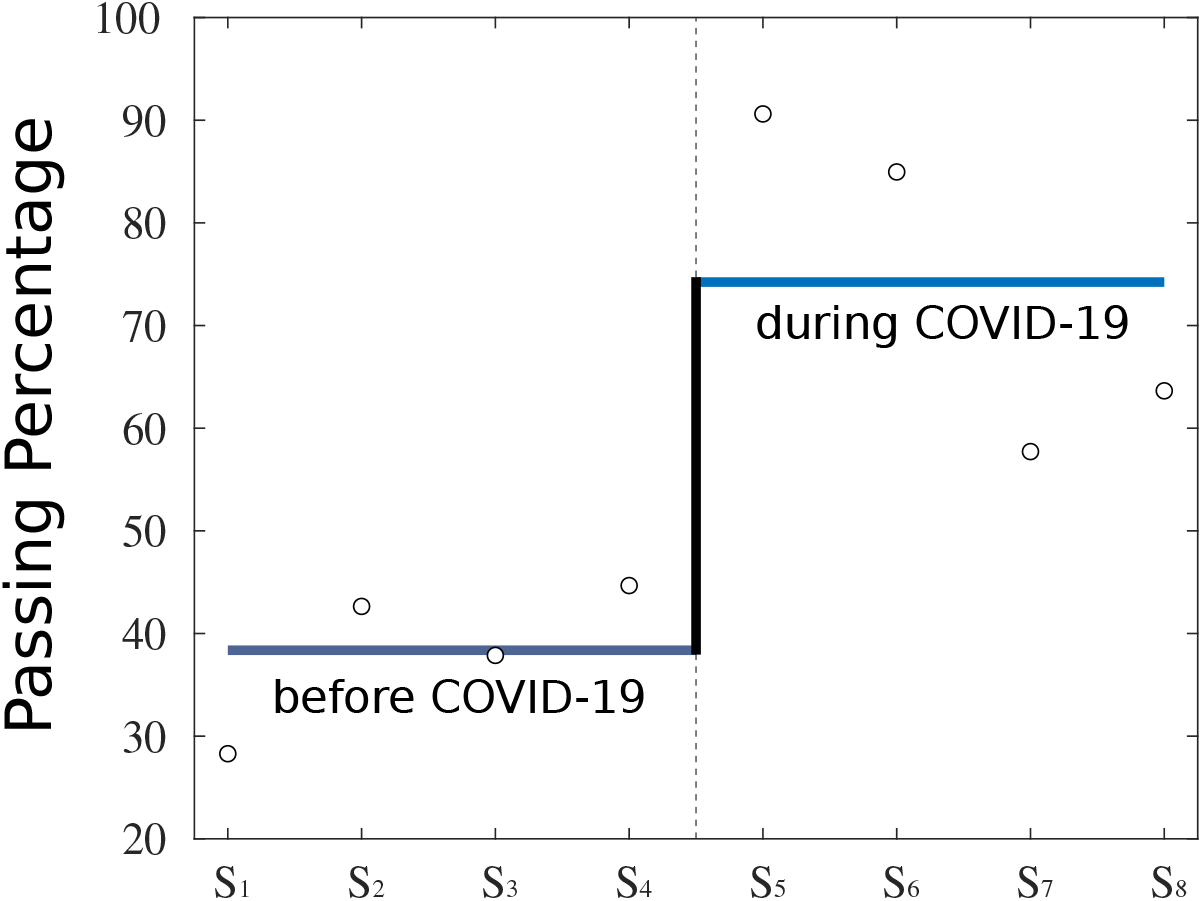
Passing percentage for Course X, four semesters before and during the COVID-19 pandemic. Except from the students of this study, during the COVID-19 pandemic, around 800 students each semester were evaluated using entirely v-tests. The passing percentage for each semester is presented with circles and its average with continue blue lines. The passing percentage shows a clear transition and an average improvement of 35.87%. Although a similar behavior can be observed in every first year course. The difference of these mean values or jumps depends on the type of evaluation applied and in the teaching resources used in each course. For a course evaluated entirely with p-tests, this improvement during the pandemic was only of 6%.

A wrong lecture of these results could lead to the conclusion that education improved during the pandemic, however in this paper we have looked inside the data, and questioned the reasons of this improvement. Whether a higher passing percentage is related with a quality education where the student is learning more and better, or not, is left as an open question.

## 4. Discussion

Virtual education is not the future. If something, was a forced reality for which no one was prepared. During the pandemic, teachers and students had to face not only virtual education but also confinement, anxiety, depression, illness and death, and any solution adopted (not only in education) was a small and improvised patch. We had to adapt abruptly to a new reality looking for new ways to teach and to learn, possibly with the best intentions. This made teachers to try methods which in other circumstances would never have been applied and surely, the major concern was the evaluations (given the problems we already had before the pandemic), resulting in the improvised experiment of the virtual assessments. On the other hand, maybe students in their desperation and frustration for feeling they were not learning much, look for a collective collaboration in order to overcome the experience. This, without necessarily meaning they were learning but at least, that they were passing.

However, the human being adapts, and there is the possibility these methods, evaluations and behaviors continue after the pandemic, just because we got used and because now results more comfortable. Mistakes were made which profoundly damaged the education in its very foundations and they have to be admitted, faced, but more importantly corrected. A great gap appeared between the ones who could learn despite the circumstances, and the ones who could not because of the circumstances. A self-criticism and analysis of the education during the pandemic is needed, not only to fill this gap, but also looking in the future, not to mention that virtual education has visualized the flaws that surely were and are present in online and remote programs.

The most important question is what did we learn from this experience? and what did we **really** learn? as teachers, students, and society. There are many things we got from virtual education, for example, the use of technological resources and the availability of the recorded material after the lectures. However virtual education and online programs will never fulfill one of the main components of education which is the human interaction and the university is nurtured with experiences, relationships, and friendship, and not only with courses which have to be passed. Maybe virtual lectures are not the actual problem as we all are used to learn from YouTube tutorials. The problem could be the method used to transmit the information, as there are very few good tutorials which use slides. Every assigned activity should be reviewed and the student should receive feedback for his work which should be original. However, during the pandemic, the student was bombed by activities which were graded but not checked at all. These plenty pendent pointless activities and a bad virtual lecture forced the student in the direction of the collective cooperation. Apart from that: is the student really learning by doing all these activities?, can these activities and v-tests really replace a traditional exam? and more important, can we guarantee these activities and evaluations are done by the student by himself? However, if the answer to the last question is no, then the first ones are unnecessary as all the educative process loses its sense. The v-tests applied during the pandemic seems that not only favored academic dishonesty but also corrupted honest students. The collaboration for cheating in a exam would be then, a tragic answer to the question: What did we really learn?

## Acknowledgments

E.G. thanks the Institution which participated in this study for its cooperation and the data provided. Special thanks to those who were my students during the COVID-19 pandemic who share their experiences and concerns, and to my dog Camila who kept me company during the virtual lectures.

